# ILRUN downregulates ACE2 expression and blocks infection of human cells by SARS-CoV-2

**DOI:** 10.1101/2020.11.13.381343

**Authors:** Leon Tribolet, Marina R. Alexander, Aaron M. Brice, Petrus Jansen van Vuren, Christina L. Rootes, Kostlend Mara, Meg McDonald, Kerri L. Bruce, Tamara J. Gough, Shuning Shi, Christopher Cowled, Andrew G. D. Bean, Cameron R. Stewart

## Abstract

The human protein-coding gene ILRUN (inflammation and lipid regulator with UBA-like and NBR1-like domain, previously C6orf106) is a recently-characterised inhibitor of the transcription regulators p300 and CREB-binding protein (CBP). Here we have utilised RNA-seq to define cellular pathways regulated by ILRUN in the context of severe acute respiratory syndrome-associated coronavirus-2 (SARS-CoV-2) infection. We find that inhibition of ILRUN expression increases cellular expression of several members of the renin-angiotensin aldosterone system (RAAS), including the SARS-CoV-2 entry receptor angiotensin converting enzyme 2 (ACE2). Furthermore, inhibition of ILRUN results in increased SARS-CoV-2 replication. These data identify ILRUN as a novel inhibitor of SARS-CoV-2 replication and represents, to our knowledge, the first report of ILRUN as a regulator of the RAAS.

**SIGNIFICANCE STATEMENT:** There is no doubt that the current rapid global spread of COVID-19 has had significant and far-reaching impacts on our health and economy and will continue to do so. Research in emerging infectious diseases, such as severe acute respiratory syndrome-associated coronavirus (SARS-CoV-2), is growing rapidly, with new breakthroughs in the understanding of host-virus interactions and the development of innovative and exciting therapeutic strategies and new knowledge and tools to better protect against the impacts of disease. The human protein-coding gene ILRUN is a recently-characterised inhibitor of the transcription regulators p300 and CREB-binding protein (CBP). Here we present the first evidence that ILRUN modulation has implications for SARS-CoV-2 infections. Virus infectivity assays confirmed that gene silencing of ILRUN had a proviral effect and increased SARS-CoV-2 replication, whilst over-expression of ILRUN inhibited SARS-CoV-2 production. Additionally, we observed that ILRUN also regulates the expression of key elements of the RAAS. These data have important implications for the development of antiviral strategies to deal with the current SARS-CoV-2 pandemic.

## INTRODUCTION

Severe acute respiratory syndrome coronavirus-2 (SARS-CoV-2, family *Coronaviridae*) is the causative agent of the current coronavirus disease (COVID-19) pandemic, one of the deadliest infectious disease events to have occurred in recent times. Since emerging in Wuhan, Hubei province, China in December 2019 (1), SARS-CoV-2 has spread across the globe, causing an estimated (as of 7 August 2020) 27 million infections and over 880,000 deaths (2). Comorbidities feature in a high percentage of deaths with hypertension being the foremost, followed by diabetes and cardiovascular disease (3). There are currently no FDA-approved medicines or therapies available to treat COVID-19, with most mitigation strategies focused on physical distancing and other public health measures (4).

The development of medical countermeasures against COVID-19 is guided by knowledge of cellular factors associated with SARS-CoV-2 infection. Viruses are obligate intracellular pathogens that rely on the hijacking of cellular machinery for successful completion of the infection cycle. SARS-CoV-2 entry requires the type-I membrane-bound glycoprotein angiotensin-converting enzyme 2 (ACE2) (5). As its name suggests, ACE2 normally functions in the renin-angiotensin-aldosterone system (RAAS) as a carboxypeptidase enzyme that converts angiotensin (Ang)-I and Ang-II into smaller peptides (Ang1-9 and Ang1-7, respectively), counteracting their vasoconstrictive properties and protecting the host from hypertension and severe cardiac dysfunction (6). The ACE2 protein is largely expressed in lung alveolar epithelial cells consistent with prominent lung pathology in SARS-CoV-2 infection (7). Other permissive cell-types predicted by protein-proofed transcript profiling include cardiomyocytes and intestinal enterocytes, consistent with heart injury and the less common intestinal symptoms (8, 9). The surface-exposed SARS-CoV-2 Spike (S) glycoprotein binds ACE2 to facilitate viral attachment to target cells, while the host serine protease transmembrane protease serine 2 (TMPRSS2) cleaves the S protein to facilitate receptor recognition and membrane fusion (10). Clinical trials are currently examining RAAS modulators, including the Ang receptor blocker Losartan in COVID-19 (11), based on the hypothesis that SARS-CoV-2 impairs the regulatory function of ACE2 and leads to unopposed RAAS activation and tissue damage. Such studies highlight how the study of host pathways associated with SARS-CoV-2 pathogenesis can facilitate the rapid clinical testing of novel countermeasures.

The human protein-coding gene ILRUN (inflammation and lipid regulator with UBA-like and NBR1-like domains, previously C6orf106) was recently characterised as a novel inhibitor of the transcription regulators p300 and CREB-binding protein (CBP) (12). In the context of virus infection, ILRUN promotes the degradation of p300/CBP to impair the function of the transcription factor interferon (IFN)-regulatory factor 3 (IRF3), thereby acting as a negative regulator of the expression of antiviral (type I IFNs) and pro-inflammatory (tumor necrosis factor, TNF) cytokines (12). ILRUN is highly phylogenetically conserved and associated with several disease states (including cancer, coronary artery disease and obesity) (13), suggesting it may possess biological functions beyond regulation the host antiviral response.

To gain further insight into ILRUN function and explore its potential roles in COVID-19 pathogenicity, we performed a transcriptomics study to identify host pathways regulated by ILRUN in the context of SARS-CoV-2 infection. RNA-seq was utilized to quantify both host and viral transcripts regulated by ILRUN in human epithelial colorectal adenocarcinoma (Caco-2) cells. Importantly in the context of SARS-CoV-2 infection, ILRUN gene knockdown leads to upregulated expression of multiple RAAS pathway members, including ACE2. In addition, we show that ILRUN expression negatively correlates with SARS-CoV-2 infection. These data suggest that ILRUN functions as a negative regulator of the RAAS pathway and is antiviral towards SARS-CoV-2. To our knowledge this study represents the first comprehensive analysis of cellular pathways transcriptionally regulated by ILRUN and suggests that ILRUN is a major checkpoint of both SARS-CoV-2 infection and the RAAS.

## RESULTS

### ILRUN function and SARS-CoV-2 infection in Caco-2 cells

To investigate host pathways regulated by ILRUN in the context of SARS-CoV-2 infection, we firstly established an *in vitro* system where cells were receptive to both RNAi-mediated gene silencing and SARS-CoV-2 infection. Previous studies have shown that Caco-2 (human colon epithelial) and Calu-3 (human lung epithelial) cells support SARS-CoV-2 infection (14). Both these cell types are derived from tissues shown to be productively infected during SARS-CoV-2 infection and contribute to disease symptoms (15, 16). siRNA reagents targeting ILRUN resulted in >75 % decrease in Caco-2 ILRUN mRNA expression compared to cells transfected with siNEG, a negative control siRNA (Fig 1A). To assess that ILRUN function in Caco-2 cells, cells were treated with transfected poly(I:C), a dsRNA mimetic that is recognised by TLR3, retinoic acid-inducible gene-I (RIG-I) and melanoma differentiation-associated gene 5 (MDA5) that acts as viral RNA mimetic that induces a type I IFN response regulated by ILRUN (12). Stimulating Caco-2 cells with poly(I:C) resulted in a robust induction of *IFNb, TNF* and *IL6* expression, which was heightened in siILRUN cells (Fig 1B), thereby validating ILRUN inhibition of the type-I IFN response in Caco-2 cells. Despite several rounds of optimisation experiments, Calu-3 cells were found to be unreceptive to siRNA or cDNA transfection in our hands (data not shown) and were not investigated further in this study.

**Fig 1.**
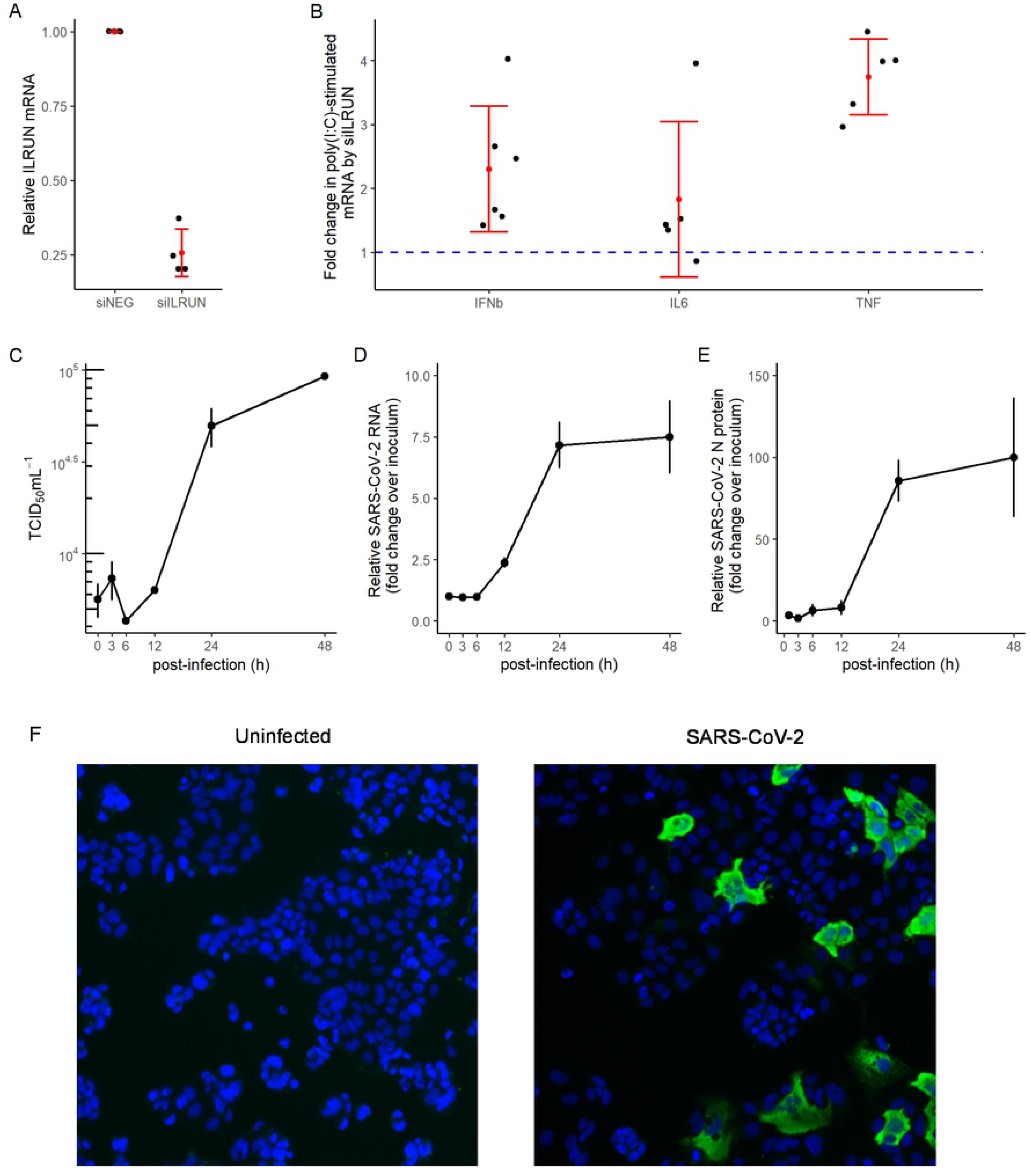
Validation of ILRUN function and SARS-CoV-2 infection in Caco-2 cells. (A) *ILRUN* mRNA levels (2^−ΔΔCt^ relative to *GAPDH*) in Caco-2 cells transfected with siRNAs (40 nM, 72 h) targeting *ILRUN* or a nontargeting control (siNEG). **p<0.01 (B) Caco-2 cells were transfected with siRNAs (40 nM, 72 h) and then treated with transfected poly(I:C) (10 μg/mL, 6 h). Cells were then harvested and analyzed for mRNA expression of the listed cytokines by qRT-PCR, normalised to *GAPDH* *p<0.05, **p<0.01. (C) TCID_50_ measurements of virus titres, (D) qRT-PCR measurements of intracellular viral RNA represented by 2^−ΔΔCt^ normalised first to *GAPDH* and then to inoculum levels of SARS-CoV-2, set to 1, and (E) intracellular viral protein in Caco-2 cells infected with SARS-CoV-2 (MOI 0.1). *p<0.05 compared to inoculum. (F) Immunofluorescence microscopy showing SARS-CoV-2 N protein staining (green) in Caco-2 cells infected with SARS-CoV-2 (MOI 0.1, 24 h). Cell nuclei were stained using DAPI (blue).

We next performed a time course experiment to characterize the single-cycle replication kinetics of SARS-CoV-2 in Caco-2 cells. A previous study has shown that increases in SARS-CoV-2 levels in tissue culture supernatant of infected Caco-2 cells are observed within 24 h post-infection (h.p.i.) (14). Consistent with this report, our experiments showed that Caco-2 cells infected with SARS-CoV-2 started producing infectious virions (above inoculum levels) between 12 and 24 h.p.i. (Fig 1C). This indicates that the length of one cycle of SARS-CoV-2 infection in Caco-2 cells is approximately 12 to 24 h. These results were validated using a qRT-PCR assay (17) to measure intracellular viral RNA levels at 3, 6, 12, 24 and 48 h.p.i.. Intracellular viral RNA levels started increasing above inoculum levels between 6 and 12 h.p.i. (Fig 1D), which is consistent with the replication kinetics observed with the virus production data (Fig 1C). When measured by quantitative fluorescence microscopy, intracellular viral nucleoprotein (N) protein levels were also observed to increase between 6 and 12 h, maximising at 24 h.p.i. (Fig 1E & F). We did not observe SARS-CoV-2 infection to induce syncytia or other cytopathic effects in Caco-2 cells (Fig 1F).

### ILRUN regulates multiple cellular pathways, including the renin-angiotensin aldosterone system

Using Caco-2 cells as a model system, we next utilised RNA-seq to identify genes differentially transcribed as a result of reduced ILRUN expression. 100bp single end Illumina sequencing reads from two Nova Seq 6000 lanes (technical replicates) were mapped to 33,121 annotated human genes. Using the DESeq2 R package to normalize read counts for each gene we found that silencing of ILRUN results in a dramatic change in the Caco-2 transcriptome. Unsupervised analysis of variance using principle component analysis (PCA) showed tight clustering of treatment groups (technical replicates not shown for clarity) (Fig 2A). By comparing siNEG to siILRUN at 72 h post transfection we found 457 differentially expressed genes using a conservative cut-off of p-value <0.05, log_2_ fold change (FC) >1 and baseMean >5 (Fig 2B and Supplementary Table 1). *ILRUN* itself was identified as differentially expressed in this analysis, validating both siRNA transfection and data analysis protocols. Many of the genes regulated by ILRUN encode proteins with proven roles in innate immunity, including Toll-like receptor 7 (*TLR7*), interferon-stimulated gene *(ISG)15,* vimentin (*VIM*) and interleukin −6 receptor (*IL6R*). Differential expression of a subset of these genes, including *ACE2*, *ISG15, VIM and IL6R* were validated by qRT-PCR (Fig 2C).

**Fig 2.**
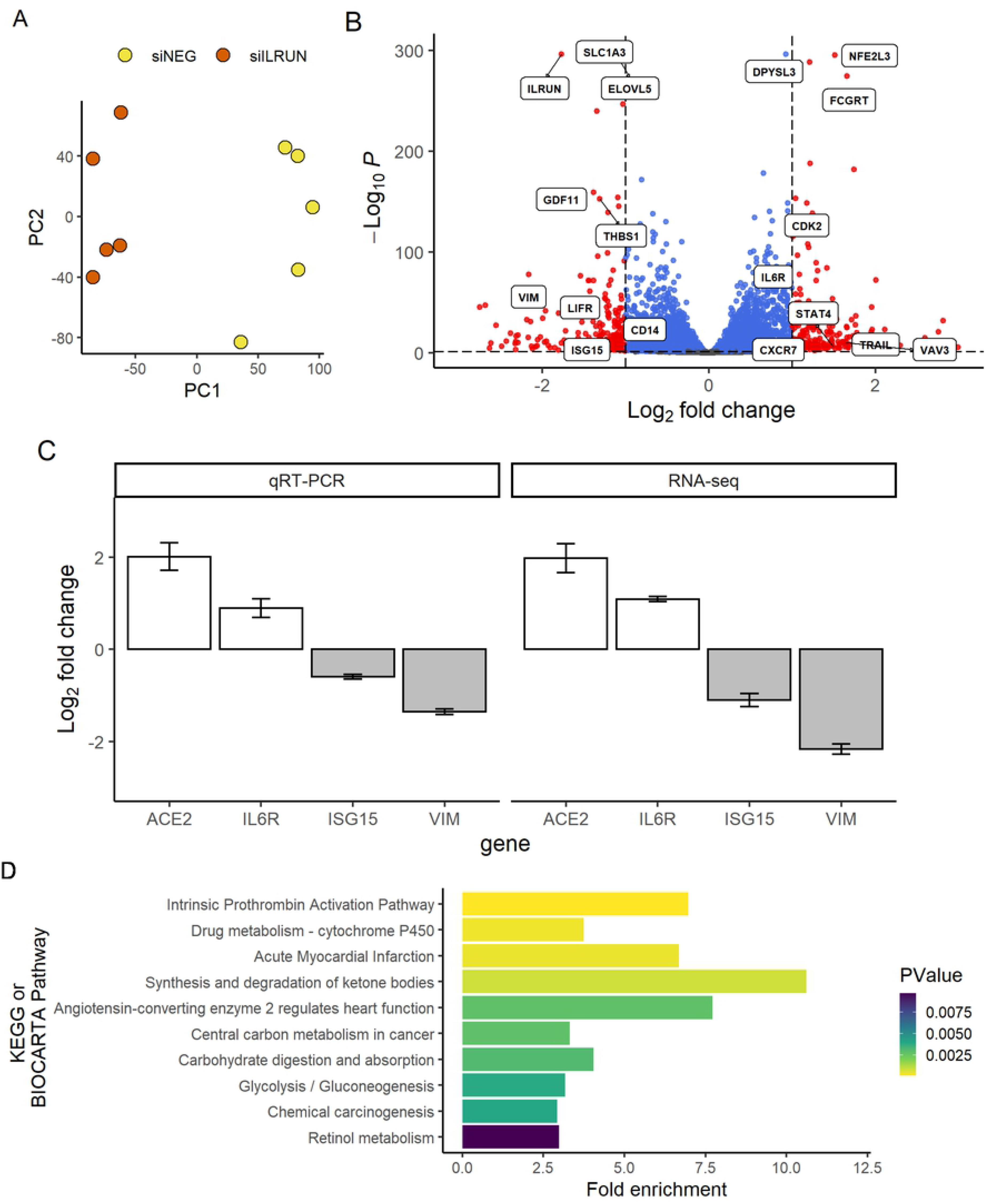
Host pathways regulated by ILRUN in Caco-2 cells. (A) Unsupervised variance analysis of 5 replicate samples transfected with 40 nM siNEG or siILRUN for 72 h. Samples plotted based on principle component (PC) 1 and PC 2 dimensions. (B) Volcano plot showing global transcriptional changes in Caco-2 cells following siRNA transfection (40 nM, 72 h). 457 transcripts (shown in red) were differentially expressed by ILRUN knockdown based on the cut-off of p-value <0.05, log2fold change (FC) >1 and baseMean >5. Genes that did not exceed the fold-change threshold are shown in blue. Genes of interest related to antiviral immunity are labelled. (C) Validation of RNA-seq differential expression by qRT-PCR for *ISG15*, *IL6R* and *VIM* in Caco-2 cells transfected with siRNAs (40 nM, 72 h). qRT-PCR differential expression was calculated based on relative expression and normalized to *GAPDH*. *p <0.05, *\*\** p <.01. (D) Functional Annotation Clustering of 457 differentially-expressed genes in host pathways (BIOCARTA and KEGG) are plotted against the fold enrichment and ordered on p-value.

Next, functional annotation clustering was performed to determine whether particular molecular pathways were overrepresented by the set of ILRUN-regulated genes. The significantly enriched pathways were largely associated with metabolism, inflammation and coagulation. Importantly, these include pathways that are distinct from those previously established to be regulated by ILRUN, namely the type-I IFN system. Of particular interest was the enrichment of the ACE2 regulates heart function pathway as ACE2 is the cell entry receptor of SARS-CoV-2. Indeed, our transcriptomics revealed that ACE2 was significantly upregulated following ILRUN knockdown, along with several other genes of the RAAS, the specific pathway in which ACE2 carries out its physiological function, including *ACE, AGT, AP-A and AP-N* (Fig 3). Together, our analysis has revealed that ILRUN likely participates in numerous biological pathways with diverse roles in cell biology, including as a novel regulator of the RAAS.

**Fig 3.**
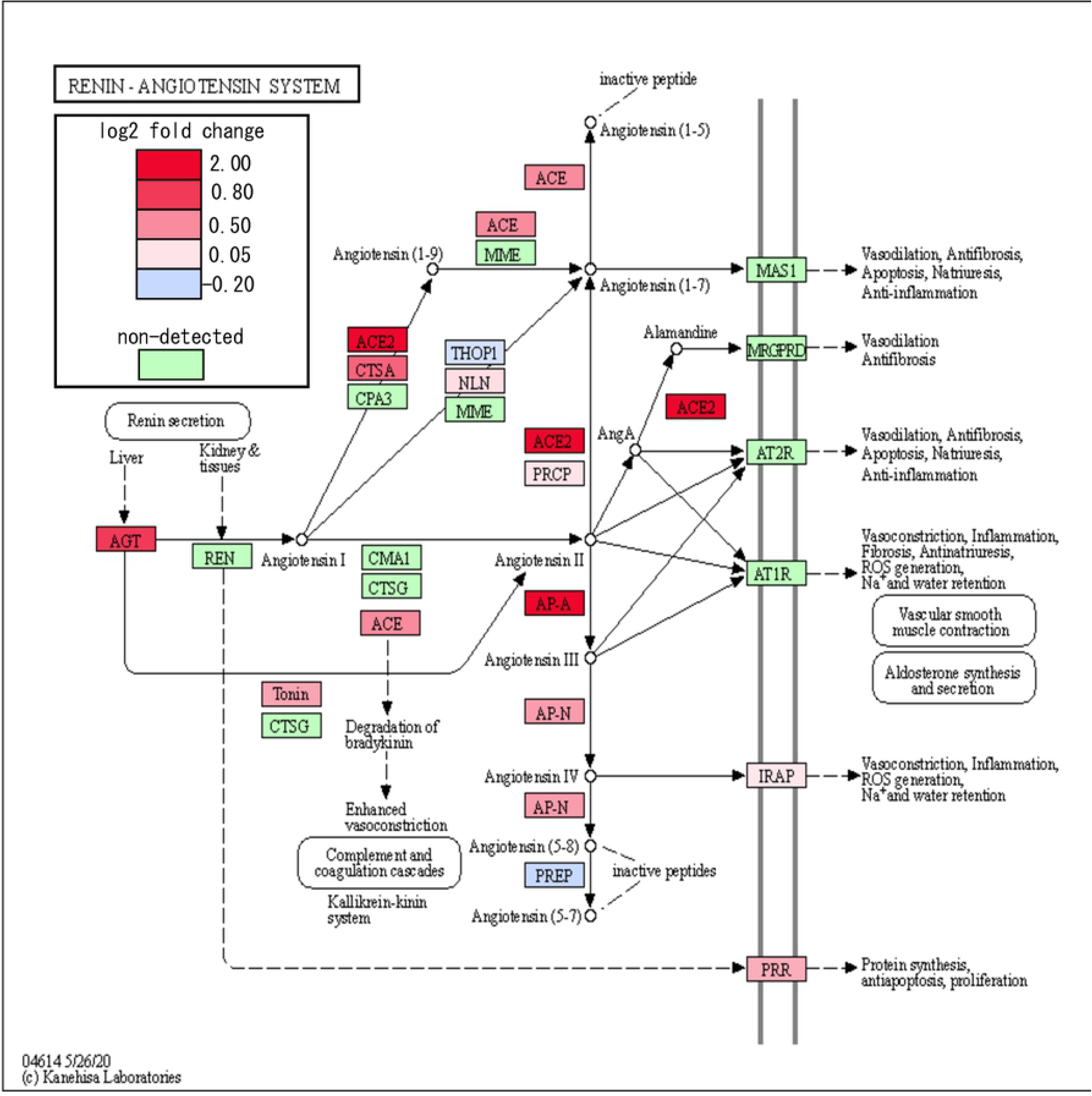
ILRUN regulates expression of key RAAS genes. The impact on siILRUN on the RAAS was investigated by quantitatively colouring the KEGG pathway map using a diverging colour palette to show fold-change effects across the entire pathway (not just those genes we had already identified as DE). Shades of red indicate up-regulation while shades of blue indicate down-regulation. Genes that were not detected in this study are shown in green.

### ILRUN suppresses SARS-CoV-2 growth by regulating the expression of enzymes critical to cell entry

Due to its regulation of ACE2 and its previous association with virus infection (18), we next examined the impact of modulating ILRUN expression on SARS-CoV-2 infection. RNA-seq reads from siNEG and siILRUN cells infected with SARS-CoV-2 for 6 h and 24 h were mapped to the SARS-CoV-2 genome using Bowtie2. After normalising for library size, we detected significantly elevated reads mapping to SARS-CoV-2 in cells depleted of ILRUN (Fig 4A). Virus infectivity assays confirmed that siILRUN increased SARS-CoV-2 production in Caco-2 cells, whilst over-expression of ILRUN inhibited SARS-CoV-2 production (Fig 4B).

**Fig 4.**
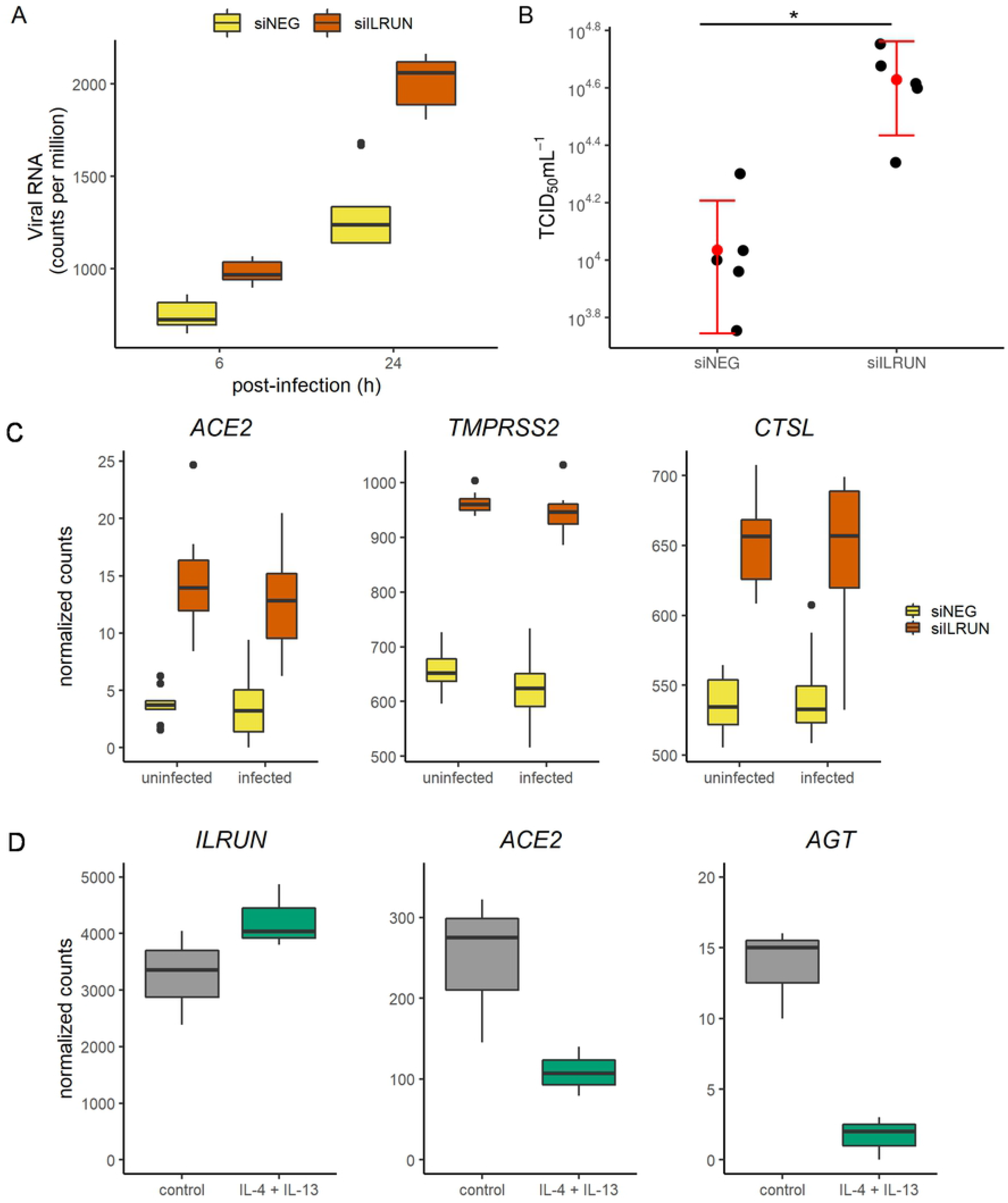
ILRUN supresses SARS-CoV-2 infection and down-regulates host genes essential for SARS-CoV-2 entry. (A) Transcription profile of SARS-CoV-2 in Caco-2 cells transfected with 40 nM siNEG or siILRUN for 72 h at 6 h and 24 h post infection. (B) SARS-CoV-2 titres of supernatants from Caco-2 cells infected with SARS-CoV-2 (24 h, MOI 0.3) post-transfection with siRNAs (40 nM, 72 h) *p<0.05. (C) Normalised RNA-seq counts measuring expression of *ACE2*, *TMPRSS2* and *CTSL* in Caco-2 cells transfected with siRNAs (40 nM, 72 h) and infected with SARS-CoV-2 (24 h, MOI 0.1) (D) *ILRUN*, *ACE2* and *AGT* expression in primary human bronchial epithelial cells stimulated with Th2 cytokines (30 ng/mL of IL-4 and IL-13) for 24 h <u>GSE113185</u>.

The antiviral properties of ILRUN against SARS-CoV-2 infection prompted further analysis of ILRUN-regulated host genes linked to SARS-CoV-2 infection. Referring to a list of host genes required by SARS-COV-2 in a recent CRISPR screen in VEROE6 cells (https://www.ncbi.nlm.nih.gov/pmc/articles/PMC7457610/), we found that three of these genes were differentially expressed upon ILRUN knockdown, namely ACE2, CTSL, and TMPRSS2. While ACE2 facilitates the attachment of SARS-CoV-2 to target cells, the proteases TMPRSS2 and cathepsin L (*CTSL*) are required for SARS-CoV-2 virus-host cell fusion (19, 20). Expression levels of all three genes were significantly upregulated in cells with reduced ILRUN, both in the absence and presence of SARS-CoV-2 (Fig 4C). Together, these data indicate that ILRUN has antiviral function towards SARS-CoV-2, by regulating the expression of key SARS-CoV-2 entry and fusion molecules ACE2, TMPRSS2 and CTSL.

Finally, to investigate the relationship between ILRUN and RAAS gene expression in primary cells, we accessed publicly-available RNA-seq data measuring gene expression in normal primary human bronchial epithelial cells from a single donor grown in triplicate (21). In this dataset, once fully differentiated, cells were stimulated with IL-4 and IL-13 to ellicit the Th2-type inflammation (22). Similar to our results observed in Caco-2 cells with higher levels of ILRUN (siNEG), Th2-induced increases in *ILRUN* in primary bronchial epithelial cells were concomitant with decreases in *ACE2* and *AGT* expression (Fig 4D). This observation extends the differential expression of RAAS genes by ILRUN into primary airway-derived cells which are more relevant to the lung pathology seen in COVID-19.

## DISCUSSION

Here we present the first evidence that ILRUN modulates SARS-CoV-2 infection. Virus infectivity assays confirmed that gene silencing of ILRUN increased SARS-CoV-2 replication in Caco-2 cells, whilst over-expression inhibited SARS-CoV-2 replication. Additionally, we observed that ILRUN regulates the expression of key elements of the RAAS, the body’s mechanism for regulating blood pressure and fluid homeostasis (23). Consistent with ACE2 serving as the entry receptor for SARS-CoV-2 (5, 19) (in addition to the other human coronaviruses HCoV-NL63 (24) and SARS-CoV (25)), reducing ILRUN expression promotes SARS-CoV-2 infection in Caco-2 cells.

Our data suggest that ILRUN inhibits SARS-CoV-2 infection by blocking the early stages of the virus infection cycle. Firstly, inhibition of ILRUN results in increases in intracellular viral RNA levels at 6 h post infection (Fig 4A), a timepoint that precedes virus replication in Caco-2 cells (Fig 1D). Secondly, we have shown ILRUN negatively regulates expression of three host proteins critical for coronavirus virus entry – ACE2, TMPRSS2 and cathepsin L (Fig 4C, Supplementary Table 1). While the coronavirus S protein mediates target cell attachment via ACE2 (5, 19), the virus-host membrane fusion reaction requires S protein subunits S1 and S2 to undergo a series of conformational changes induced by proteolysis, which is mediated at the plasma membrane by TMPRSS2 (19) and within endosomal compartments by cathepsin L (20). In addition to facilitating virus entry, TMPRSS2 also mediates cell-cell fusion of infected cells (26). Pharmacological inhibitors of both TMPRSS2 (19) and CTSL (20) potently inhibit coronavirus infection *in vitro*, highlighting the potential of targeting these proteases as therapeutic countermeasures against SARS-CoV-2.

Despite being highly evolutionarily conserved, with orthologues expressed in nearly all metazoans (13), ILRUN remains a relatively poorly characterised protein. Our previous studies show that ILRUN inhibits IRF3-dependent antiviral cytokine transcription, while other studies have identified *ILRUN* single nucleotide polymorphisms (SNPs) are associated with increased risk of obesity (27, 28), coronary artery disease (CAD) (29), and altered timing of pubescent growth spurts (30, 31). Our current hypothesis is that ILRUN is linked to these processes, and the RAAS, through its regulation of the histone acetyltransferases p300 and CBP. p300/CBP are considered master regulators of gene transcription that control proliferation, differentiation, infection and apoptosis, among other processes (32). *AGT* expression is tightly regulated by p300/CBP (33), while the *ACE2* promoter region also contains a p300/CBP binding site (34), nominating a potential mechanism by which ILRUN regulates expression of these genes. Notably, the RAAS is a key pathway associated with CAD, with inhibitors standardly prescribed for treatment (35). RAAS genes are also found to be upregulated in the dendritic cells of CAD patients, potentially contributing to pathogenesis. Similarly, the RAAS is involved in lipid metabolism and upregulated in adipocytes as a result of obesity (36, 37). Thus, the potential role of ILRUN in CAD and obesity may be via its transcription regulation of RAAS components.

IFNs are key to antiviral immunity and viral recognition elicits IFN production, which triggers the transcription of IFN-stimulated genes (ISGs), which engage in various antiviral activities. Nevertheless, type I IFNs are widely expressed and can result in immunopathology during viral infections. An intriguing finding from this study is that while ILRUN regulates the type I IFN induction pathway, SARS-CoV-2 did not induce a noticeable type I IFN response in Caco-2 cells, indicating this is independent of the ILRUN-induced IFN expression. Another study has similarly observed a lack of type I IFN induction in Caco-2 cells infected with SARS-CoV-2, including at a high MOI (38). While one explanation for this result is that Caco-2 cells do not mount robust innate immune response to viral infection, previous studies have shown infection of Caco-2 cells with Sendai virus and Newcastle Disease virus induces a strong type I IFN response, while SARS-CoV-1 suppresses these cytokines (39). Our own data also shows that Caco-2 cells respond to the viral RNA mimetic poly(I:C) (Fig. 1B). In Calu-3 cells (which support SARS-CoV-2 replication to higher titres than Caco-2 cells (14), SARS-CoV-2 induces a type-I IFN response that is markedly delayed compared to other viral infections such as influenza A virus (40, 41) and Sendai virus (42). SARS-CoV-2 appears to be particularly adept at immune evasion, encoding seven proteins that inhibit IFN-β promoter activation (43), among them the ORF6 protein that blocks both type I IFN induction and signalling pathways. Several groups have hypothesized that the lack of timely and robust antiviral response to SARS-CoV-2 infection contributes to COVID-19 pathogenesis (38, 43).

Hadjadj et al. showed that peripheral blood immune cells from severe and critical COVID-19 patients show enhanced inflammatory responses (44). In our analysis of note were the inflammatory associated cytokines, IL-6R and IL-1β which were differentially regulated upon suppression of ILRUN. Inflammation is known to play a crucial role in the pathogenesis of severe infections and ARDS and evidence is emerging that the IL-1/IL-6 pathway is highly upregulated in patients with severe disease. Both IL-6R and IL-1β have been shown to be significantly upregulated in severe COVID infection (44). Importantly, controlling excessive immune activation is critical as particularly as these cytokines can drive of inflammasome activation and in an autoinflammatory loop can lead to uncontrolled inflammation.

Our data also reveals that ILRUN promotes basal transcription of ISG15, a host member of the ubiquitin family that is targeted by SARS-CoV-2 as an evasion mechanism against anti-viral immune responses (45). ISG15 plays diverse roles in antiviral immunity as both a promoter and negative feedback inhibitor of IFN signalling, in addition to direct antiviral activity (46). These functions are carried out via conjugation onto both cellular and viral proteins via an enzymatic cascade (termed ISGlyation) or secreted as an unconjugated protein that acts as a cytokine (46). Coronaviruses, including SARS-CoV-2, encode papain-like protease (PLpro) that, in addition to processing viral proteins to activate the virus replicase complex, cleave post-translational modifications on host proteins to deactivate immune signally cascades (46). While the PLpro of SARS-CoV predominantly targets ubiquitin chains, SARS-CoV-2 PLpro preferentially cleaves ISG15, including from IRF3, resulting in attenuated type-I IFN signalling during infection (45). GRL-0617, a chemical compound targeting SARS-CoV-2 PLpro, restores ISG15-mediated activation of IRF3 during SARS-CoV-2 infection and inhibits virus replication (45). Beyond antiviral signalling, extracellular signal-related kinase-1 (ERK1) has been shown to be ISGylated, though the functional relevance of this modification has yet to be elucidated (46). ERK1 participates in the Ras-Raf-MEK-ERK MAP kinases signalling pathway, which regulate the cell-cycle in response to extracellular signals (47). Notably, ILRUN has been shown to activate a member of this pathway, 90 kDa ribosomal r6 kinase (p90RSK), a target of ERK1 (48). The presence of a UBA-like domain in ILRUN (13), which generally facilitate binding to ubiquitin and ubiquitin-like protein, also offers the intriguing possibility that ILRUN may directly interact with ISG15 to modulate its function. Thus, future studies to elucidate the functions and mode of action of ILRUN may additionally further our understanding of the complex roles of ISG15 in cellular biology, including during infection with pathogenic viruses such as SARS-CoV-2.

In summary, this study sheds further light on the regulatory function of ILRUN, a recently characterised inhibitor of p300/CBP with links to infectious disease, cancer, obesity and cardiovascular disease. The study reveals a previously unappreciated role for ILRUN in the regulation of the RAAS and, remarkably, three key genes (*ACE2*, *TMPRSS2* and *CTSL*) required for infection of human cells by SARS-CoV-2 and other coronaviruses. Future work will aim to understand the molecular mechanisms underlying ILRUN function and progress this molecule as a potential drug target for a number of diseases of significant concern to human health.

## MATERIALS AND METHODS

### Cell lines

VeroE6 cells (ATCC CRL-1586) were maintained in Gibco Dulbecco’s Modified Eagles Medium (DMEM) supplemented with 10 % (v/v) foetal calf serum (FCS), 100 U/mL penicillin, and 100 μg/mL streptomycin (Life Technologies). Caco-2 cells were maintained in Gibco Modified Eagles Medium (MEM) supplemented with 20 % (v/v) FCS, 10 mM HEPES, 0.1 mM non-essential amino acids, 2 mM glutamine, 1 mM sodium pyruvate, 100 U/mL penicillin, and 100 μg/mL streptomycin (Life Technologies). All cells were kept at 37 °C in a humidified incubator (5% CO_2_).

### Virus

All virology work was conducted at the CSIRO Australian Centre for Disease Preparedness at physical containment (PC)-4. The isolate of SARS-CoV-2 (BetaCoV/Australia/VIC01/2020) was received from the Victorian Infectious Disease Reference Laboratory (VIDRL, Melbourne, Australia) and passaged in VeroE6 cells for isolation, followed by passaging in VeroE6 cells for stock generation. All virus stocks were aliquoted and stored at −80°C for inoculations.

### Transfections

Caco-2 cells were transfected with 40 nM small interfering (si)RNA (GE Life Sciences) and 2 μL (24-well plates) or 0.6 μl (96-well plates) Dharmafect-1 (GE Life Sciences) in serum-free Minimum Essential Medium Eagle (MEM, Life Technologies). For DNA transfections plasmids encoding ILRUN-FLAG (pILRUN) and green fluorescent protein (GFP) are described in (12), 100 ng DNA was incubated with 0.4 μL FugeneHD (Promega) in serum-free MEM. Cells were stimulated with transfected high molecular weight poly(I:C) (Invivogen) (10 μg/mL with 3 μL FugeneHD) for 6 h. *RNA-seq*

Total RNA was isolated from cells in Trizol using the RNeasy kit (Qiagen) as per the manufacturer’s instructions. The quality and quantity of RNA was assessed for all samples using a Bioanalyzer (Agilent, Santa Carla, USA). RNA-Seq was performed by the Australian Genome Research Facility (AGRF). Illumina TruSeq Stranded mRNA libraries were prepared, followed by sequencing on an Illumina Novaseq-6000 for generation of 100 base pair (bp) single-end reads on two lanes (technical replicates). Between ~15-21 million raw single-end reads were obtained per sample. Raw data were assessed for overall quality using fastqc v0.11.8. (http://www.bioinformatics.babraham.ac.uk/projects/fastqc/).

### Bioinformatic analysis of RNA-seq data

Quality and adapter trimming was performed using TrimGalore v0.6.4 (http://www.bioinformatics.babraham.ac.uk/projects/trim_galore/) with default settings for automatic adapter detection. Trimmed reads were mapped to the National Cener for Biotechnology Information (NCBI) human reference genome (GRCh38) downloaded from Illumina iGenomes (http://igenomes.illumina.com.s3-website-us-east-1.amazonaws.com/Homo_sapiens/NCBI/GRCh38/Homo_sapiens_NCBI_GRCh38.ta r.gz) using Tophat v2.1.1 (49). The number of reads overlapping each gene in the NCBI annotated reference genome (GRCh38) were counted using htseq-count v0.11.2 within Python v3.7.2 (50), using the intersection-nonempty mode to handle reads overlapping more than one feature. A custom python script was used to compile counts from individual samples into a single count matrix file. The Bioconductor package DESeq2 was used to test for differential expression between different experimental groups (51). Volcano plots were prepared in R using (https://github.com/kevinblighe/EnhancedVolcano). Gene lists were submitted to DAVID bioinformatics resources (https://david.ncifcrf.gov/summary.jsp) and Functional Annotation Clustering performed using default settings to retrieve GO terms in each annotation cluster. For Kyoto Encyclopedia of Genes and Genomes (KEGG) pathway mapping (52), custom python scripts were used to prepare input lists for conversion of gene symbols to Entrez gene IDs using bioDBnet’s db2db tool (https://biodbnet-abcc.ncifcrf.gov/db/db2db.php) and generation of hex codes based on fold change for colour mapping of KEGG pathways (https://www.genome.jp/kegg/tool/map_pathway2.html). Trimmed reads were mapped to a SARS-CoV-2 reference genome of the virus used in experiments (MT007544; SARS-CoV-2/human/AUS/VIC01/2020) using bowtie v2.3.4 (53). Mapped viral reads were counted using samtools v1.10.0 (Li et al., 2009) and coverview v1.4.4 (54) then normalised to library read content. Normalized counts from viral reads and DESeq2 (host genes) were plotted using geom_boxplot function in the ggplot2 R package using the 25^th^ and 75^th^ percentiles to form the box and whiskers no larger than 1.5 times the interquartile range. Data points beyond the whiskers are outliers.

### Analysis of publicly available data

The NCBI Gene Expression Omnibus was searched for suitable RNA-seq datasets and GSE Accession number entered into the GREIN : GEO RNA-seq Experiments Interactive Navigator (http://www.ilincs.org/apps/grein/) to retrieve metadata and normalized counts. Normalised counts were plotted for each gene using ggplot2 package in R.

### Quantitative real-time PCR

500 ng of RNA was reverse transcribed to DNA using SensiFast reverse transcriptase (Bioline) first-strand cDNA synthesis protocols. qRT-PCR was performed on a Quant Studio 3 thermocycle (Applied Biosystems) with Taqman probes for SARS-CoV-2 E gene, *ILRUN* (Hs00256056_s1), *IFNβ* (Hs01077958_s1)*, TNFα* (Hs00174128_m1)*, IL6* (Hs00174131_m1)*, ISG15* (Hs01921423_s1), *IL6R* (Hs01075664_m1), *VIM* (Hs00958111_m1) and glyceraldeyde-3-phosphate dehydrogenase (*GAPDH)* (Hs02758991_g1) were purchased from ThermoFisher Scientific. ACE2 mRNA expression was measured using SYBR Green dye and primers (F: GGAGTTGTGATGGGAGTGAT, R: GATGGAGGCATAAGGATTTT) as described (55). PCR cycling for gene detection was at 95 °C for 10 min followed by 45 cycles of 95 °C for 15 sec and 60 °C for 1 min. RNA transcription data were analysed using the 2^−ΔΔCT^ method and were normalized to GAPDH expression.

### TCID_50_ analysis

The infectious titres of SARS-CoV-2 stocks was determined by TCID_50_ assays performed as described previously (56). Samples were titrated in quadruplicate in 96-well plates, co-cultured with VeroE6 cells for four days and monitored for development of cytopathic effects (CPE). Infectious titres were calculated by the method of Reed and Muench (57).

### Immunofluorescence and quantification of relative antigen staining

Caco-2 cells were fixed for 30 min in 4 % paraformaldehyde (PFA) and stained with a polyclonal antibody targeting the SARS-CoV-2 Nucleocapsid (N) protein (Sino Biological, catalogue number: 40588-T62, used at 1/2,000) for 1 h. Cells were subsequently stained with 1/1,000 dilution of an anti-rabbit AF488 antibody (Invitrogen catalogue number A11008). Nuclei were counter-stained with diamidino-2-phenylindole (DAPI). Caco-2 cells were imaged using the CellInsight quantitative fluorescence microscope (Thermo Fisher Scientific) at a magnification of 10 x, 49 fields/well, capturing the entire well. The relative viral antigen staining was quantified using the Compartmental analysis bioapplication of the Cellomics Scan software.

### *Statistic*s

The difference between two groups was analysed in GraphPad Prism by a two-tailed Student’s *t* test and between multiple groups by one-way ANOVA. A *P* value of <0.05 was considered significant. All data points are the average of biological triplicates, with error bars representing standard error of the mean. All data are representative of results from at least 2 separate experiments.

## ABBREVIATIONS

ILRUN: inflammation and lipid regulator with UBA-like and NBR1-like domains
ACE: angiotensin converting enzyme
CBP: CREB binding protein
COVID-19: coronavirus disease 2019
SARS-COV-2: severe acute respiratory syndrome coronavirus 2
RAAS: renin angiotensin aldosterone system
Ang: angiotensin
IFN: interferon
TNF: tumor necrosis factor
siRNA: short interfering RNA
IL-: interleukin
qRT-PCR: quantitative real-time polymerase chain reaction
h.p.i: hours post-infection
N: nucleocapsid
TLR: Toll-like receptor
VIM: Vimentin
CTSL: cathepsin L
TMPRSS2: transmembrane protease serine 2
MOI: multiplicity of infection
PLpro, ISG: interferon-stimulated genes
ARB: angiotensin receptor blocker
ACEI: angiotensin converting enzyme inhibitor
FCS: foetal calf serum

## AUTHOR CONTACT INFORMATION

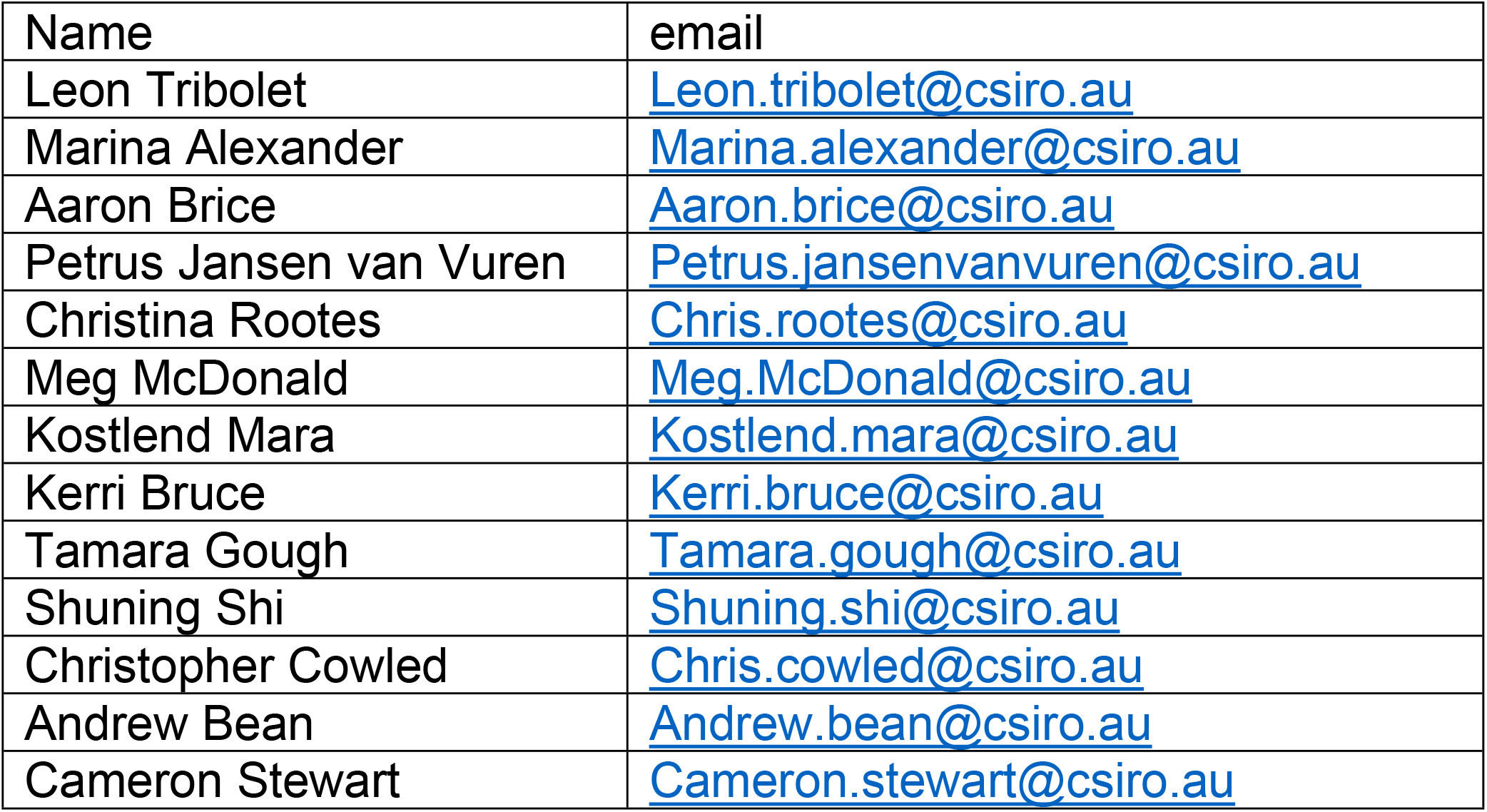

The mailing address for all authors is:

CSIRO Australian Centre for Disease Preparedness

Private bag 24

Geelong 3220 Victoria

Australia

